# Noise Analysis in Biochemical Complex Formation

**DOI:** 10.1101/310847

**Authors:** Zikai Xu, Khem Raj Ghusinga, Abhyudai Singh

## Abstract

Several biological functions are carried out via complexes that are formed via multimerization of either a single species (homomers) or multiple species (heteromers). Given functional relevance of these complexes, it is arguably desired to maintain their level at a set point and minimize fluctuations around it. Here we consider two simple models of complex formation – one for homomer and another for heteromer of two species – and analyze how important model parameters affect the noise in complex level. In particular, we study effects of (i) sensitivity of the complex formation rate with respect to constituting species’ abundance, and (ii) relative stability of the complex as compared with that of the constituents. By employing an approximate moment analysis, we find that for a given steady state level, there is an optimal sensitivity that minimizes noise (quantified by fano-factor; variance/mean) in the complex level. Furthermore, the noise becomes smaller if the complex is less stable than its constituents. Finally, for the heteromer case, our findings show that noise is enhanced if the complex is comparatively more sensitive to one constituent. We briefly discuss implications of our result for general complex formation processes.

## 1 Introduction

Living cells comprise of numerous molecular species that interact with each other. A widely present theme across these interactions is the assembly of a macromolecule or complex from a number of constituents. Such complexes may be present both as homomers (i.e., made from a single species) and heteromers (made from multiple species). For example, the protein holin in bacteriophage λ’s lysis system is known to form multimer with itself, as well as with its counterpart anti-holin [1, 2]. Among many other examples of biochemical complexes are signaling molecules [3–7], ribosomes [8, 9] and other macromolecular machineries [10, 11], etc.

Formation of biochemical complexes plays vital role for most cellular processes, including gene regulation, signal transduction [3–7]. Given their importance, it can be argued that their level must be closely regulated so as to achieve robust function. However, many species are present at low copy numbers in cells, and thereby stochasticity (or noise) in the reactions involving them is unavoidable [12]. Previous work indeed has shown that complex formation might be an important mechanism in control of noise in gene regulation [13–15].

In this paper, we investigate how various attributes of the complex formation process affect the noise (quantified as fano factor, variance/mean) in the complex level. We focus on the steady-state noise in two toy models of complex formation. In the first model, a single species forms a homomer whereas in the second model two species interact to form a heteromer. Not only these toy models themselves are appropriate for analysis of some real biological examples [16], but they also hint towards what behaviors might arise in more involved scenarios.

Our strategy to analyze the noise behavior relies upon using moment dynamics of these toy models. Due to nonlinearities in present in them, however, the moment dynamics is not closed and we use a linear approximation of the system at its steady-state to estimate the noise behavior. To this end, we first carry out deterministic analysis of the reaction systems to find their fixed points. Under appropriate assumptions on the complex formation rates for both models, we show that they exhibit unique real fixed points. We then linearize the complex formation rates around their respective fixed points and write moment dynamics of the approximate systems.

We focus on how two key parameters affect the noise in steady state complex level: (i) sensitivity of the complex formation rate with respect to the species’ level, and (ii) relative stability of the complex in comparison with that of the constituting species. Interestingly, our analysis reveals that noise in complex level has a *U*-shape profile with respect to sensitivity for both homomer and heteromers. Moreover, we find that for these toy models, if the complex is relatively unstable (i.e., it degrades faster) as compared with its constituents, the overall noise profile shifts downwards. We also analyze the heteromer with different sensitivities for each species. In this case, the overall noise profile shifts upwards.

The paper is organized as follows. Section 2 discusses both homomer and heteromer models and provides deterministic analysis to find their fixed points. In Section 3, the stochastic description of these models and corresponding moment dynamics are presented. The moment dynamics is linearized around the fixed points that we obtained in Section 2. Section 4 explores the effect of sensitivity, and relative stability on the noise, and finally Section 5 concludes the paper.

*Notation:* We use bold letters to represent stochastic processes, such as *X*; whereas non-bold letters correspond to deterministic processes, such as *x*. Expectation operator is denoted by 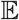. The set of real numbers and natural numbers are respectively denoted by 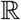 and 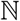. We also use 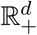 to denote positive real numbers in a *d*-dimensional Euclidean space. To make physical sense, species counts and rates of various reactions are taken to be positive.

## 2 Stochastic model formulation

In this section, we describe two simple models of complex formation (Fig. 1). The first model consists of a single species that forms a homomer, whereas the second model consists of two species that combine to form a heteromer.

**Figure. 1:**
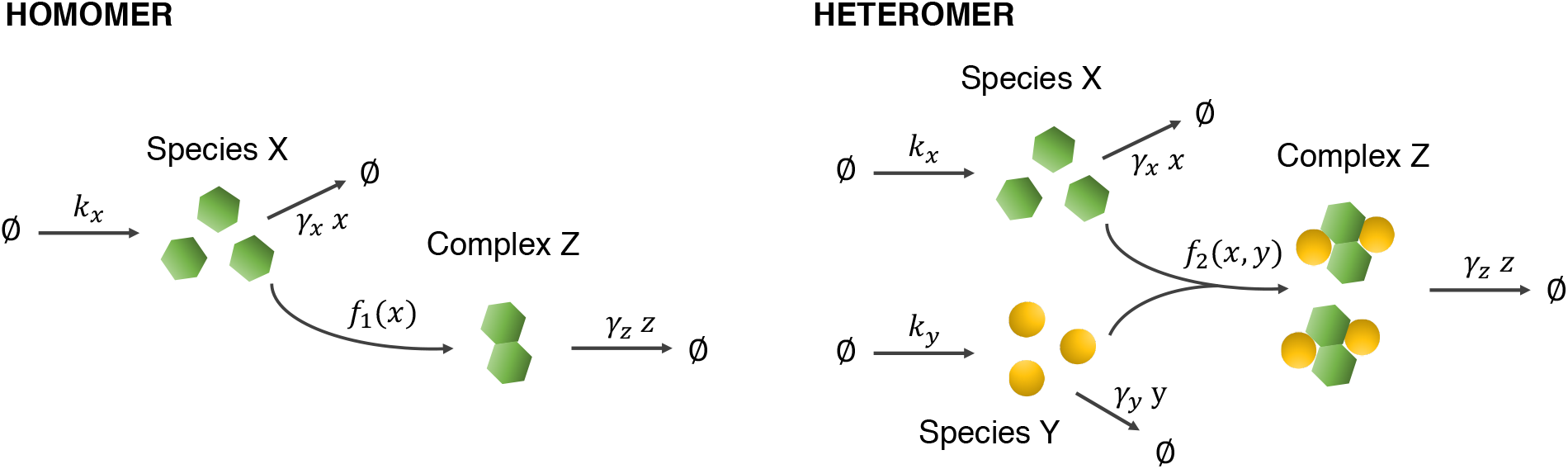
Schematic for complex formation. *Top:* A species *X* is produced at a rate *k_x_* and its *n_x_* molecules combine to form a complex (homomer), *Z*, with rate *f*_1_(*x*). The species *X* and the complex *Z* degrade enzymatically with rates *γ_x_x* and *γ_z_z* respectively. *Bottom:* Two species, *X* and *Y*, are respectively produced at rates *k_x_* and *k_y_. n_x_* molecules of *X* combine with *n_y_* molecules of *Y* to form a complex (heteromer), *Z*, with rate *f*_2_(*x*, *y*). Both constituting species and the complex decay enzymatically with corresponding rates *γ_x_x, γ_y_y* and *γ_z_Z*.

### 2.1 Homomer

Consider the following biochemical system that comprises of production and degradation of a species, *X*. Furthermore, *n_x_* molecules of *X* interact to form a complex *Z*, which can also degrade.

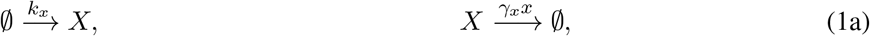

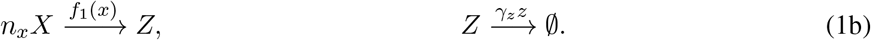

Here the levels of species *X* and the complex *Z* are denoted by small letters *x* and *z* respectively. The production rates of *X* follows a zero-order kinetics with rate constant *k_x_* while the complex *Z* is made with a rate *f*_1_(*x*) where *f*_1_ denotes an arbitrary positive function in its argument. Assuming a mass action kinetics, *f*_1_(*x*) would take the form *k*_1_*x^n_x_^* where *k*_1_ > 0; however, other forms of *f*_1_(*x*) are also possible, e.g., Michelis-Menten, Hill Function, etc. While we do not assume a particular form of *f*_1_ we assume that it satisfies the following properties:

a. lim_*x*→0+_ *f*_1_(*x*) = 0
b. *f*_1_(*x*) is a monotonically increasing function in *x*.

These assumptions are made to ensure that the system in (1) has unique positive steady state. Finally, degradation of both *X*, and *Z* is assumed to follow first-order reaction kinetics with rates *γ_x_*, and *γ_z_*.

One way to describe the dynamics of the biochemical system in (1) is to use the ordinary differential equation (ODE) based approach. In this case, the following ODEs can be used to compute (*x*, *z*)

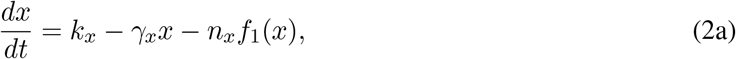

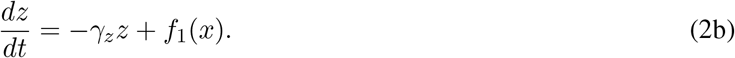

With the aforementioned assumptions on *f*_1_ (*x*), it can be shown that 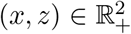 is positively invariant for the dynamical system in (2). That is, if the system starts from an initial condition in 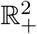, then it remains in that set. In addition, (2) has a unique steady-state in 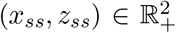 which is given by the real-valued solution to the following algebraic system

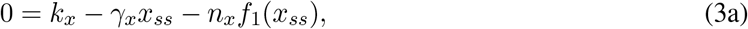

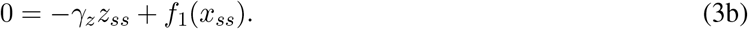

We refer interested readers to Appendix A for a sketch of proof of these statements.

### 2.2 Heteromer

Next consider the scenario where two species *X* and *Y* interact to form the complex *Z*. The biochemical system now consists of production of *X* and *Y*, interaction between them to form *Z*, and degradation of all three of them

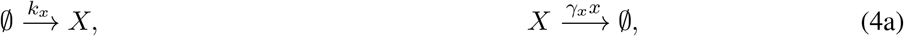

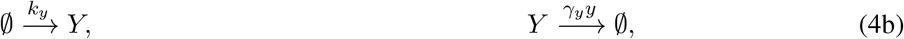

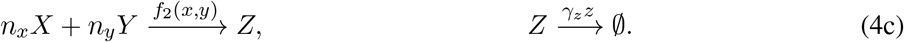

Here, the notations *x* and *z* have same meaning as the homomer case, and likewise *y* denotes the level of *Y*. The production rates of *X* and *Y* are *k_x_* and *k_y_* while the complex *Z* is made with a rate *f*_2_(*x*, *y*) where *f*_2_ denotes an arbitrary positive function of its arguments. As in the homomer case, we do not assume a particular form of *f*_2_. However in order to ensure that the system in (4) has unique positive steady state, we assume that it satisfies the following properties

1. lim_*x*→0+_ *f*_2_(*x*, *y*) = 0,
2. lim_*y*→0+_ *f*_2_(*x*, *y*) = 0,
3. *f*_2_(*x*, *y*) is a monotonically increasing function in both its arguments.

Lastly, each of the species *X*, *Y* and *Z* are assumed to degrade enzymatically with rates *γ_x_*, *γ_y_* and *γ_z_* respectively.

In a deterministic description, the time evolution of (*x*, *y*, *z*) is governed by the following system of differential equations

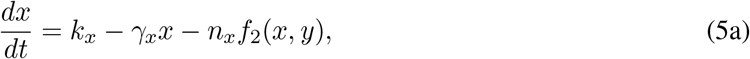

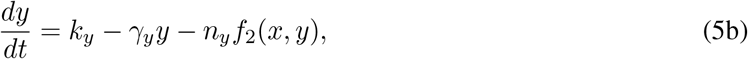

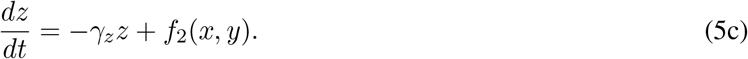

With the previously mentioned assumptions on *f*_2_(*x*, *y*), it can be seen that (5) has a unique steady-state in 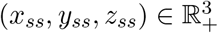 which is given by the real-valued solution to the algebraic system

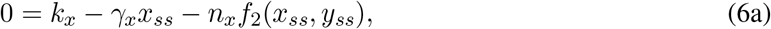

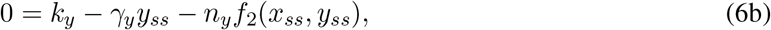

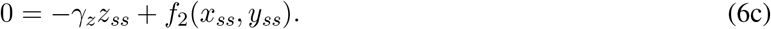

Sketch of the proof for these properties is provided in Appendix A.

While the above deterministic descriptions of homomer/heteromer can provide insights into some important system behaviors (e.g., response times), they fail in accounting for stochastic effects. The stochasticity naturally arises from the probabilistic nature of reactions, but it becomes particularly prominent when the reacting species are present in low copy numbers [17]. Our focus in this work is to analyze the stochastic behavior of the system, particularly the noise in complex level. To this end, we provide stochastic descriptions of the systems in (1) and (4) in the subsequent section.

## 3 Stochastic Models and Moment Equations

Let x(*t*), y(*t*) and z(*t*) respectively denote the molecular counts of *X*, *Y* and *Z* (the species *Y* is only applicable for the heteromer case). While x(*t*), y(*t*), z(*t*) are stochastic processes, we will omit the explicit dependence on time unless the context demands it. The time evolution of these can be described via probabilities of various events happening in an infinitesimal time-interval (*t*, *t* + *dt*]. We describe the detailed stochastic models for both cases in Table 1.

**Table 1:** Description of Stochastic Models for Complex Formation

| Model | Event | Population Reset Function | Propensity |
| --- | --- | --- | --- |
| **Homomer** | Production of $X$ | $\mathbf{x}(t) \mapsto \mathbf{x}(t) + 1$ | $k_x$ |
| | Degradation of $X$ | $\mathbf{x}(t) \mapsto \mathbf{x}(t) - 1$ | $\gamma_x \mathbf{x}(t)$ |
| | Production of $Z$ | $\mathbf{z}(t) \mapsto \mathbf{z}(t) + 1$ | $f_1(\mathbf{x})$ |
| | | $\mathbf{x}(t) \mapsto \mathbf{x}(t) - n_x$ | $f_1(\mathbf{x})$ |
| | Degradation of $Z$ | $\mathbf{z}(t) \mapsto \mathbf{z}(t) - 1$ | $\gamma_z \mathbf{z}(t)$ |
| **Heteromer** | Production of $X$ | $\mathbf{x}(t) \mapsto \mathbf{x}(t) + 1$ | $k_x$ |
| | Degradation of $X$ | $\mathbf{x}(t) \mapsto \mathbf{x}(t) - 1$ | $\gamma_x \mathbf{x}(t)$ |
| | Production of $Y$ | $\mathbf{y}(t) \mapsto \mathbf{y}(t) + 1$ | $k_y$ |
| | Degradation of $Y$ | $\mathbf{y}(t) \mapsto \mathbf{y}(t) - 1$ | $\gamma_y \mathbf{y}(t)$ |
| | Production of $Z$ | $\mathbf{z}(t) \mapsto \mathbf{z}(t) + 1$ | $f_2(\mathbf{x}, \mathbf{y})$ |
| | | $\mathbf{x}(t) \mapsto \mathbf{x}(t) - n_x$ | $f_2(\mathbf{x}, \mathbf{y})$ |
| | | $\mathbf{y}(t) \mapsto \mathbf{y}(t) - n_y$ | $f_2(\mathbf{x}, \mathbf{y})$ |
| | Degradation of $Z$ | $\mathbf{z}(t) \mapsto \mathbf{z}(t) - 1$ | $\gamma_z \mathbf{z}(t)$ |

Ideally one needs to solve the chemical master equation in order to fully characterize a stochastic model [18, 19]. However, here we are only interested in studying the noise in complex level. Therefore, we directly use the dynamical equations to compute first two moments. Below we provide moment equations for both homomer and heteromer models of the complex formation process.

### 3.1 Moment Dynamics for Homomer

Using standard tools from stochastic systems [18, 19], the time evolution of expected value of a monomial x^*m*_1_^ z^*m*_2_^, with 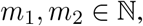, can be written as

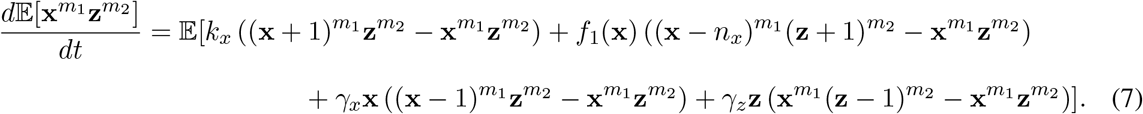

Here order of a moment is given by the sum *m*_1_ + *m*_2_.

An important point to note is that if the complex formation rate, *f*_1_ (x), is constant or a linear affine function of x, then dynamics of a moment of a certain order can be described in terms of moments of same or lower order. However, if *f*_1_ (x) is is nonlinear, then moment dynamics is not closed: dynamics of a moment depends on moments of order higher than it. This is referred to as the problem of moment closure. Several *moment closure* methods that provide approximation to moment dynamics have been proposed in literature [17, 20–31]. These methods are based on several themes such as linearization, neglecting some higher order moments/cumulants, assumptions on underlying distribution, preservation of some dynamical, physical or moment properties, etc. [20, 21]. In this paper, we use a linearization technique called linear noise approximation wherein the nonlinear propensity *f*_1_ (x) is linearized around the deterministic solution to (1) [18, 31]. This technique is known to give accurate approximations in the limit of low noise.

Recall that because of our assumptions on the function *f*_1_(*x*), the system (1) has a unique, positive, real equilibrium point. Assuming small fluctuations in (x, z) around the steady-state deterministic solution (*x_ss_*, *z_ss_*), we linearize *f*_1_(x) and substitute the linearized form in (7). More specifically, we take

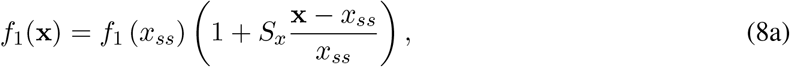

where

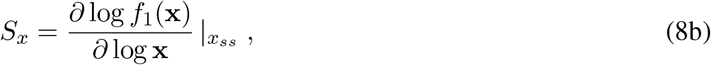

is the log sensitivity of *f*_1_(x) with respect to x. Plugging (8) in (7), the dynamics of first two moments of each of the species can be obtained. We then use these to compute approximate first two stationary moments of the complex *Z* and eventually compute the noise in *Z*.

### 3.2 Moment Dynamics for Heteromer

Next we consider the heterodimer wherein *X* and *Y* combine to form the complex *Z*. In this case, expected value of a monomial x^*m*_1_^ y^*m*_2_^ z^*m*_3_^, 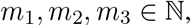, is governed by the following ODE

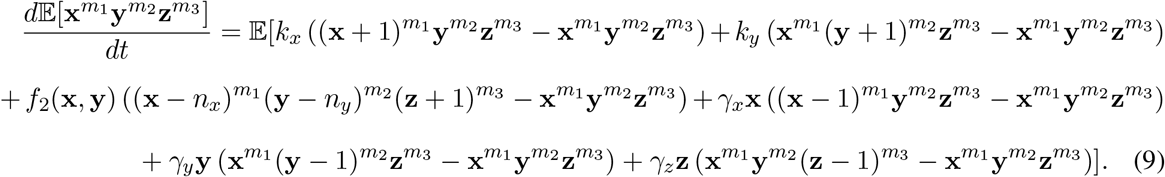

As with the homomer case, the assumptions on *f*_2_(x, y) imply that (4) has a unique, positive, real equilibrium point. We can linearize *f*_2_(x, y) around the steady-state deterministic solution (*x_ss_*, *y_ss_*, *z_ss_*) More specifically, we take

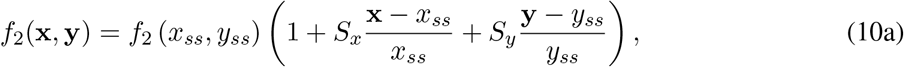

where

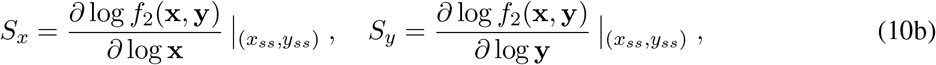

are the log sensitivities of *f*(x, y) with respect to x and y. Plugging (10) in (9), the dynamics of first two moments of each of the species can be obtained. We can use symbolic computations in Mathematica for this purpose. It turns out that the form of the second moment of *Z* is quite convoluted, which subsequently results in convoluted expression for the fano factor *FF_Z_*. We do not provide those expressions here and only provide formula for *FF_Z_* for a simple case wherein *S_x_* = *S_y_, x_ss_ = y_ss_, γ_x_ = γ_y_*, and *n_x_* = *n_y_* (see the next Section). For other cases, we have to rely on numerical computation.

## 4 Analysis of Noise Properties of the Complex

In this section, we analyze how various parameters in the complex formation process affect the noise in the complex level. For this purpose, we quantify the noise in *Z* using fano factor (variance/mean). We investigate the effect of varying the sensitivity with respect to one of the constituents and relative degradation rate of the complex as compared with those of its constituents.

### 4.1 Steady-State Noise in Homomer level

To systematically analyze the effect of important model parameters on the noise, we keep the steady-state means in the linearized model constant (note that they are exactly same as the deterministic steady-state solution to (1)). More specifically, we find the production rate of *X* in terms of other model parameters and fixed steady-state means (*x_ss_, z_ss_*)

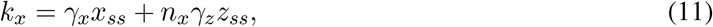

Using this we obtain the following expression for the fano factor (*FF_Z_*)

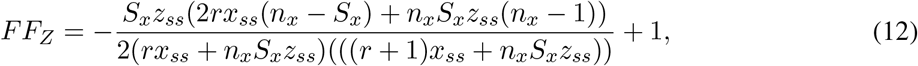

where *r* = *γ_x_/γ_z_* is the ratio of degradation rates of the species *X* and the complex *Z*.

Analyzing the above formula provides several important insights in to how noise is affected by various parameters. For example, *FF_Z_* varies non-monotonically with respect to sensitivity *S_x_*. When the sensitivity is small, production rate of *Z* is approximately constant. Indeed *FF_Z_* takes a value close to one, which corresponds to a Poisson limit (Fig. 2, Top). With increase in *S_x_*, the noise first decreases and increases back after an optimal value. Further, if the number of molecules required to form the homomer, i.e., *n_x_*, is increased then the overall noise profile shifts downwards, which suggests that a homomer made of more molecules is better in terms of noise suppression.

**Figure. 2:**
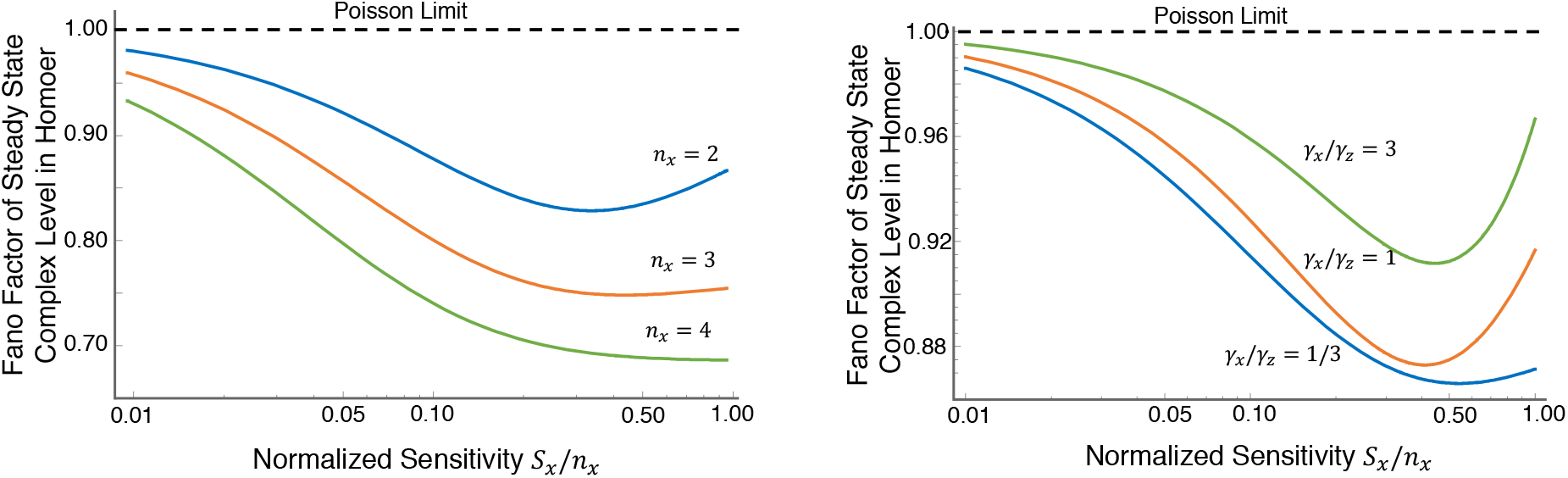
Noise in complex level as a function of sensitivity of complex formation rate with respect to the species, and relative degradation rates of the complex and the species. *Top*: The noise in complex level shows a non-monotonic behavior with increase in the sensitivity (normalized by the stoichiometry). The noise further decreases when the stoichiometry of the species in the complex formation process is is higher, suggesting that a higher order multimer can suppress noise better. The noise approaches the Poisson limit of low sensitivity values. *Bottom:*. The non-monotonic curve between noise and sensitivity shifts upwards as the relative degradation rates of the species and the complex are increased. Thus, the noise increases when the complex is relatively unstable than the species.

It is also important to point out that we have treated *S_x_* and *n_x_* separately since we consider a general form of *f*_1_(*x*). However, if *f*_1_(*x*) is assumed to follow a mass-action kinetics, then it can be easily checked from (8) that *S_x_ = n_x_*. For this reason, we only consider the values of *S_x_* between 0 to *n_x_* to be physiologically relevant and accordingly scale the *S_x_* axis in Fig. 2.

In addition to *S_x_*, we also investigate how stability of the species *X* and the complex *Z* determine the noise in *Z*. To this end, we note that the fano factor in (12) only depends on the parameter *r* = *γ_x_/γ_z_*, and not on individual values of the degradation rates. Also, we can find the limit of *FF_Z_* for large/small *r* as

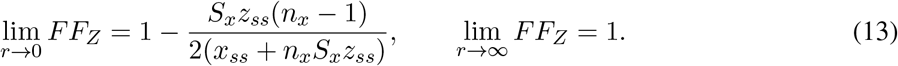

As shown in Fig. 2 (Bottom), increasing *γ_x_/γ_z_* shifts the noise vs sensitivity curve upwards.Thus, making the complex relatively unstable with respect to the species results in lower noise in the complex.

### 4.2 Steady-State Noise in Heteromer level

As in the homomer case above, we analyze the noise in the heteromer level while maintaining the steady-state of the system at some (*x_ss_, y_ss_, z_ss_*). To this end, the production rates of *X* and *Y* are varied such that the following hold

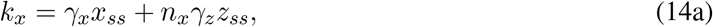

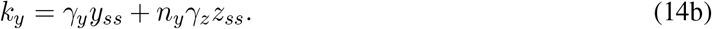

It turns out that while the expression for the fano factor *FF_Z_* is quite complicated for a general case, a simpler form can be obtained for a special case when *n_x_* = *n_y_*, *S_x_* = *S_y_* and *γ_x_* = *γ_y_*.

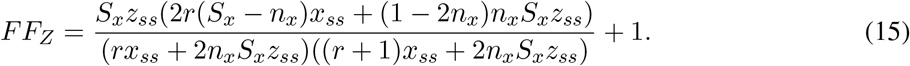

Note that in this case *x_ss_* is equal to *y_ss_*, while *r* = *γ_x_/γ_z_*. A close examination of (15) shows that it resembles (12) except for some scaling factors. Not surprisingly, even in this case varying *S_x_* produces a non-monotonic, *U*-shape behavior for noise in *Z* (we do not show the results here). We further look at how the results change when *S_x_* ≠ *S_y_*. Interestingly, it is seen that the noise always increases in this case, showing that if the complex formation rate is more sensitive to one constituent than another then its results in a higher noise in the complex level (Fig. 3).

**Figure. 3:**
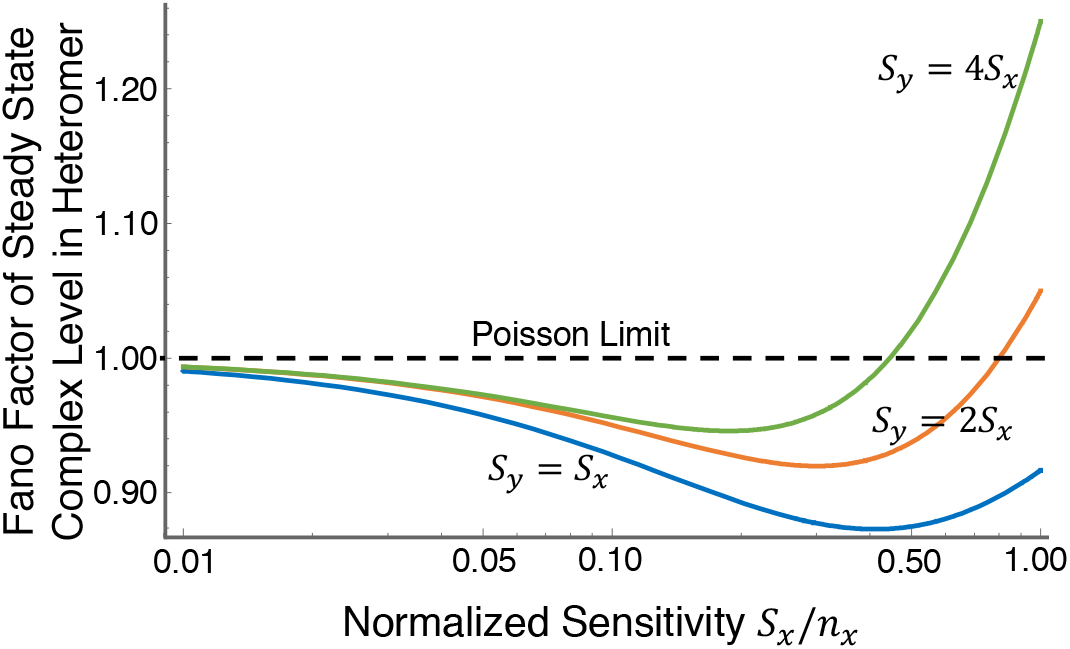
Noise in complex level when complex formation rate has different sensitivities to each constituent. While the noise behavior shows non-monotonic profile with increase in sensitivity with respect to a constituent, the curve shifts upwards as the sensitivities of both species are changed. This implies that a symmetric sensitivity is better for noise suppression.

## 5 Conclusion and Future Work

In this paper, we analyzed two simple models of biochemical complex formation. The first model was of a homomer that is formed from a single species, whereas the second model was of a heteromer of two species.

We explored the effect of two parameters in the models: (i) sensitivity of the complex formation rate to the species, and (ii) relative stability of the species and the complex. Key insights from our analysis are:

- For a homomer, its steady state noise shows a non-monotonic behavior as the sensitivity of the complex formation rate to the species level is changed. Moreover, the noise in complex level reduces when the complex is relatively unstable as compared to the species.
- For a heteromer, the noise exhibits similar behavior as in the homomer case if sensitivities and other parameters of both species are exactly same. However, when the complex formation rate is more sensitive to one species, then noise in the complex increases.

While these results are derived for simple cases, the similarity between noise behaviors of complexes made of one and two species suggests that they may hold even for complexes consisting of additional species. Since many complexes that occur in biochemical systems are made of multiple species [32], studying their noise behavior and relating it to their function would be an important direction of future research. It would also be interesting to explore reversible kinetics for the complex formation (i.e., *Z* dissociates to its constituents), and also self-regulation in production of the species as found in production of a range of proteins [33–35],

## APPENDIX

### Fixed point of (2)

We first show that for each initial condition in the positive orthant 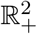, the system (2) remains in 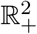. For this purpose, note that before a solution (*x*(*t*), *z*(*t*)) of (2) leaves 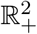 at a time *t* = *t*_1_, it has to satisfy either *x*(*t*_1_) = 0, or *z*(*t*_1_) = 0, or both. When *x*(*t*_1_) = 0, we have that 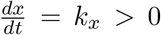 therefore, the solution cannot become 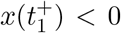. Likewise, when *z*(*t*_1_) = 0, then 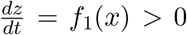. Moreover, when *x*(*t*_1_) = 0, *z*(*t*_1_) = 0, we have that 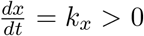 and 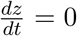. Thus, in all three cases the solution does not leave 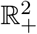.

Next, uniqueness of the fixed point can be established simply by noting that *γ_x_x* + *f*_1_(*x*) = 0 for *x* = 0 and it is an increasing function of *x*. Thus, *k_x_* and *γ_x_x* + *f*_1_ (*x*) can only be equal at one value of *x*. The solution to 0 = *k_x_* − *γ_x_x_ss_* − *f*_1_(*x_ss_*) can be then used to compute the unique value of *z_ss_* = *f*_1_(*x_ss_*)/*γ_z_*.

### Fixed point of (5)

The system in (5) also exhibits similar properties as (2), and they can be established with similar arguments as well. First we need to show that the system (5) remains in 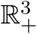 for each initial condition in 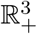. To this end, note that a solution (*x*(*t*), *y*(*t*), *z*(*t*)) of (5) can leave 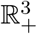 at a time *t*_1_ if one or more of *x*(*t*_1_), *y*(*t*_1_) and *z*(*t*_1_) is zero and then derivative in that direction is negative. As with the arguments for (2) in the previous section, it can be shown that it does not happen.

Next, uniqueness of the fixed point can be shown simply by noting that *γ_x_x* + *f*_2_(*x*, *y*) =0 for *x* = 0 and it is an increasing function of *x*. Similarly *γ_y_y* + *f*_2_(*x*, *y*) = 0 for *y* = 0 and it is an increasing function of *y*. Thus, for any value of *y*, *k_x_* − *γ_x_x* − *f*_2_(*x*, *y*) = 0 can only hold for one value of *x*. Likewise, for any value of *x*, *k_y_* − *γ_y_y* − *f*_2_(*x*, *y*) = 0 can only hold for one value of *y*. Collectively, one could argue that there is only one value of (x, y) for which both hold simultaneously. The solution to 0 = *k_x_ − γx_ss_ − f*_2_ (*x_ss_, y_ss_*), 0 = *k_y_ − γ_y_y_ss_ − f*_2_(*x_ss_, y_ss_*) can be then used to compute the unique value of *z_ss_* = *f*_2_(*x_ss_*, *y_ss_*)/*γ_z_*.

